# Body and brain signalling during naturalistic learning situations

**DOI:** 10.64898/2026.01.09.698657

**Authors:** Suvi Karjalainen, Juha Leukkunen, Tiina Kullberg, Tiina Parviainen

**Author notes:** Corresponding author Suvi Karjalainen Department of Psychology, Centre for Interdisciplinary Brain Research University of Jyväskylä, PO Box 35, FI-40014 Jyväskylä, Finland.

## Abstract

A central challenge in supporting learning and designing learning environments is to understand individual differences in learning experiences. However, despite growing interest in examining learning experiences in naturalistic contexts, studies employing multimethod and hyperscanning approaches remain scarce. In this study, electroencephalography (EEG) and electrocardiography (ECG) were combined to measure brain activity, heart rate, and heart rate variability simultaneously during authentic learning situations in aviation and forestry. We investigated how students’ and instructors’ cardiac and brain activity varied across different phases of a naturalistic learning situation and tasks with varying levels of difficulty. Results indicate that both students’ and instructors’ heart rate, heart rate variability, and brain activity vary systematically across behaviourally and pedagogically meaningful phases of the learning situation. These findings enhance our understanding of how physiological and neural activity vary within and across individuals during learning, providing insight into the nervous system determinants underlying individual differences in learning experiences.

## INTRODUCTION

A central challenge in supporting learning and in developing learning environments is to understand the sources of individual differences in learning and how individuality can be considered in teaching. Traditionally, individual variability in learning has been approached from different theoretical and scientific traditions ranging from cognitive to social and environmental perspectives (e.g., Chen et al., 2000; Khanolainen et al., 2020; Parhiala et al., 2018). In recent years, interest in neuroscientific and psychophysiological approaches in education research has grown and these methods are increasingly employed to investigate the functioning of the nervous system underlying individual differences in learning (e.g., Giannakos et al., 2019; Lohvansuu et al., 2021; Tomassini et al., 2011), and more recently the focus has shifted toward exploring the neurophysiological aspects of learning in naturalistic settings (Ahonen et al., 2018; Dikker et al., 2017; Silvennoinen et al., 2022; van Atteveldt et al., 2018).

From the nervous system perspective, individual differences are typically linked to structural and functional properties in the body and the brain. For example, the characteristics of rhythmic brain activity, neural oscillations, appear to reflect both intra- and inter-individual variability (Dinstein et al., 2015; Haegens et al., 2014; Pauls et al., 2024). Indeed, in addition to moment-to-moment fluctuations in brain activity (i.e., state), previous studies have shown that each individual has a relatively stable and identifiable neural “fingerprint” that distinguishes them from others (i.e., trait) (da Silva Castanheira et al., 2021; Dinstein et al., 2015; Wiesman et al., 2022). Similar to brain networks, bodily systems also operate at specific frequencies (e.g., heart rate 1.25 Hz, breathing 0.2 Hz), which vary within and across individuals (Klimesch, 2018; Nokia & Penttonen, 2022; Penttonen & Buzsáki, 2003; Soroka et al., 2025). Hence, the individual differences in learning may stem from underlying differences in the rhythmic body and brain activity.

Although it is difficult to identify any biosignal characteristics measured from the body or the brain that would directly reflect learning itself, some well-established characteristics reflecting particular learning-related processes, such as attention, vigilance, cognitive load, task engagement, excitement, and stress, can be extracted from the neurophysiological recordings. These key factors not only serve as the mechanisms that facilitate and support learning but also reflect the experiential aspects of the learning process. Interestingly, these cognitive and emotional processes are closely linked to cardiac and brain activity, including heart rate and heart rate variability (Appelhans & Luecken, 2006; Berntson & Cacioppo, 2004) as well as oscillatory brain activity (Buzsáki & Draguhn, 2004; Foxe & Snyder, 2011; Jensen & Mazaheri, 2010; Klimesch, 2012; Klimesch et al., 2007; Oken et al., 2006; Olbrich et al., 2009; Thut et al., 2012). Since these physiological and neural features vary both within and across individuals (Appelhans & Luecken, 2006; Forte et al., 2019; Haegens et al., 2014; Luque-Casado et al., 2016), examining them may shed light on individual variability in learning-related processes, and especially learning experience.

Interactive technological learning environments, such as simulations and virtual or augmented reality, offer new opportunities for investigating learning experience especially in safety-critical and repetition-intensive fields (Chernikova et al., 2020; Lateef, 2010; Petersen et al., 2022). They allow researchers to study authentic learning situations and the interactions that occur in ecologically valid contexts, while addressing methodological constraints, such as study reproducibility (van Atteveldt et al., 2018). Previous studies have shown that brain imaging and psychophysiological measures can be applied to investigate stress (Girzadas et al., 2009; Peabody et al., 2024), cognitive load (Koskelo et al., 2024; Mansikka et al., 2016; Saus et al., 2006), and emotions (Harley et al., 2019; Madsgaard et al., 2022; Vesisenaho et al., 2019) in interactive technological learning environments. The surge of such studies in the field of education, however, raises concerns about potential overinterpretation and overstated conclusions when findings are not grounded in hypothesis-driven or theory-guided experiments (Janssen et al., 2021). Despite the growing interest in applying brain imaging techniques and psychophysiological recordings to study learning within naturalistic contexts, research employing multimethod approaches and hyperscanning studies that simultaneously examine the student and the instructor remain scarce (Silvennoinen et al., 2022). The multimethod approach combined with hyperscanning used in this study offers a novel way to examine learning-related processes and learning experience in naturalistic learning situations.

In this study, we focus on well-established and interpretable measures of cardiac and brain activity, that have been previously studied in the context of learning in both laboratory and naturalistic settings. These measures not only reflect learning-related processes but also provide insight into the experiential aspects of learning. By focusing on heart rate, heart rate variability, and the characteristics of oscillatory brain activity of both students and instructors, we aim to capture how the functional state of the body and the brain vary across a naturalistic learning situation. First, we aim to determine whether robust and reliable features can be measured and extracted from cardiac and brain activity that reflect learning experience during naturalistic learning situations. Next, we aim to examine how students’ and instructors’ cardiac and brain activity vary across different phases of a natural learning situation as well as across tasks with varying levels of difficulty.

## METHODS

The present study is part of a multidisciplinary consortium project titled “Development of educational service ecosystem using physiological data and intelligent systems”. Ethical approval for this study was obtained from the local ethics committee of the University of Jyväskylä. Written informed consent was obtained from all participants before any study procedures, and the study was conducted in accordance with the Declaration of Helsinki.

### Participants

This study was conducted in naturalistic learning situations within aviation and forestry contexts. The simulation-based learning situation in the aviation context was implemented at the Finnish Air Force Academy using an in-house developed simulator, whereas in the forestry context at the Poke Vocational College, the John Deere Timberskills 4.0 simulation learning environment was used in forest machine operator training. A total of 12 student-instructor dyads in aviation (n = 6 dyads) and forestry (n = 6 dyads) participated in the study. Student-instructor dyads consisted of 12 male students (16-25 years) and 4 male simulation training instructors. In total, each instructor had three students under their guidance.

### Procedure

The data collection of each dyad lasted approximately 3.5 hours including preparations for the measurements, the simulation-based learning situation, and the debriefing. All simulation-based learning situations consisted of three phases: instructions, tasks, and feedback (Figure 1). During the simulation-based learning situation the student and the instructor were seated side by side in front of the simulator, so that they were able to interact and communicate. The instructor guided the student through the simulation.

**Figure 1.**
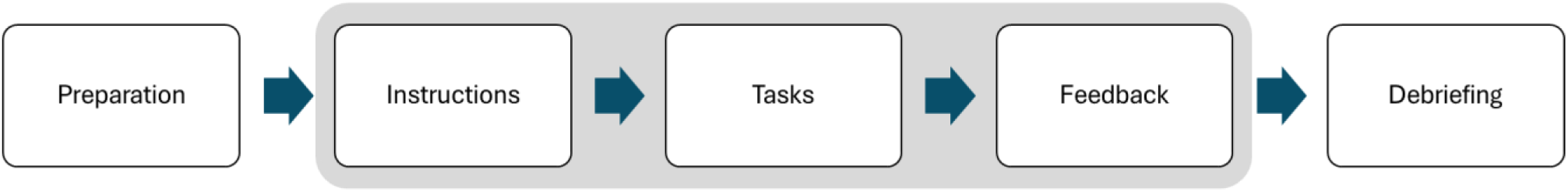
Schematic illustration of the naturalistic learning situation.

At the instructions phase, the student and the instructor went through the structure and general instructions for the upcoming tasks. More specific instructions regarding the tasks were given right before each task. During the simulation tasks, the task difficulty progressively increased so that for the final tasks students had hardly any previous experience of completing them. The tasks in the aviation context included 1) slip to the left and right, 2) pendulum, 3) barrel roll, and 4) instrument flight or pull-up during landing. In forestry context, the tasks were 1) evening log ends while loading, 2) work order while thinning, 3) stopping to a working area and thinning ready-marked trees, and 4) selecting and cutting trees for thinning. After every task, the performance of the student was briefly reviewed verbally by the instructor. After completing all the tasks, an in-depth feedback discussion was held between the student and the instructor to foster a shared understanding of the simulation-based learning experience. Subsequently, a debriefing in the form of a stimulated recall interview was conducted to obtain a comprehensive understanding of the experiences of both students and instructors during the learning situation.

### Data collection

Both neural (electroencephalogram, EEG) and cardiac activity (electrocardiogram, ECG) were recorded simultaneously during the naturalistic learning situation. EEG and ECG signals were recorded from each student-instructor dyad using the Bittium NeurOne system (Bittium Biosignals Ltd, Kuopio, Finland) with a sampling frequency of 1 kHz. EEG signals of the instructors and aviation students were recorded with a standard 64-channel EEG cap (EASYCAP, BrainProducts GmbH, Gilching, Germany) and a customized 13-channel EEG cap (channels AF3, AF4, F7, F3, F4, F8, FC5, FC6, C3, C4, CP5, CP6, Oz; Neoprene Headcap with NG Geltrode electrodes and press stud cables, Neuroelectrics, Barcelona, Spain) adapted for the VR headset was used for the forestry students. EEG caps were individually fitted and prepared for each student and instructor. To measure cardiac activity, one electrode was placed beneath each clavicle. Additionally, a ground electrode was placed on the neck and a reference electrode on the mastoid. EEG impedances and the quality of neural and cardiac signals were visually inspected and fixed if needed prior to the recording.

During the naturalistic learning situation, time stamp annotations representing the general time along with other relevant annotations related to significant events during the simulation-based learning situation, such as the beginning and end of each task, were added to the Bittium NeurOne system for later integration, synchronization, and temporal alignment of different data types. Furthermore, video recordings were used to gather detailed information on the timeline of events during the learning situation. These recordings were later reviewed to verify and refine the annotations made during the session.

### Data segmentation

To address the continuous nature of the naturalistic learning situation, the raw EEG and ECG signals were segmented based on annotations that defined the temporal progression of the distinct phases of the simulation-based learning sessions for each student-instructor dyad (Figure 2). Subsequently, these behaviourally and pedagogically meaningful, distinct phases were used for further analyses. Although the overall structure of the simulation-based learning situations was comparable across aviation and forestry contexts, slight differences were present between the aviation and forestry due to context-specific characteristics of the simulations (e.g., continuous flight with varying tasks versus discrete tasks separated by breaks). To ensure comparability, analyses focused on phases that were consistently repeated across both contexts: rest, instructions, feedback, and tasks of varying difficulty (Table 1). Finally, robust and reliable neurophysiological correlates were extracted to characterize the functional states of the body and the brain across these distinct phases of the learning situation.

**Figure 2.**
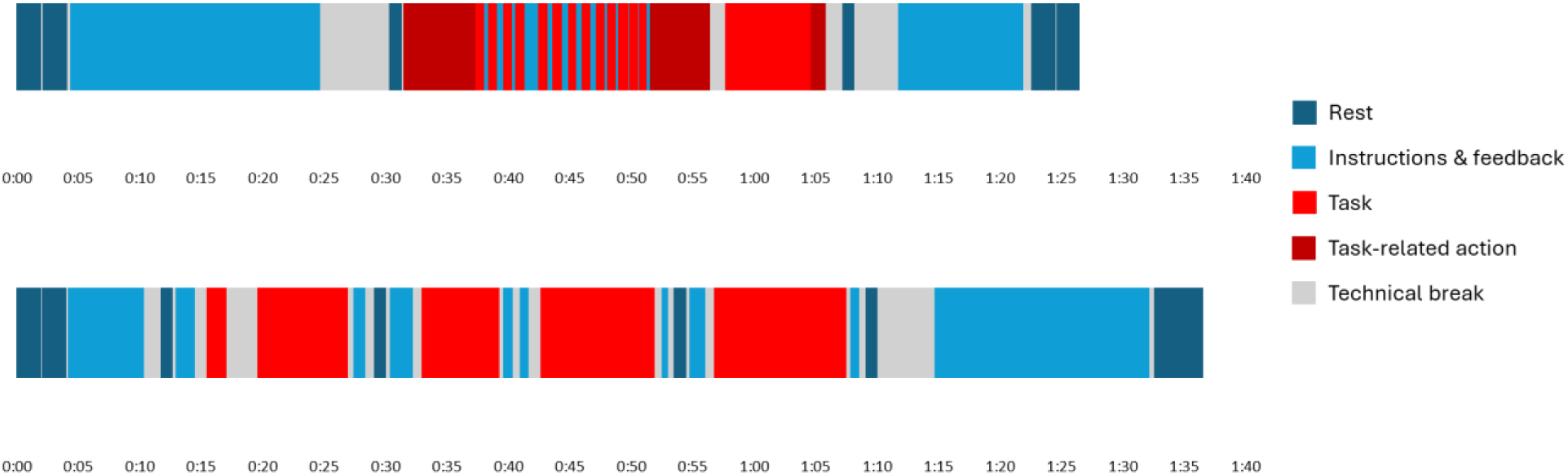
Example timelines depicting naturalistic learning situations for student-instructor dyads in aviation (upper) and forestry contexts (lower).

**Table 1.** Mean segment durations (mean ± SD, in seconds) across the different phases of the naturalistic learning situation.

| Segment | Aviation | Forestry |
| --- | --- | --- |
| Rest pre | 118 ± 2 | 120 ± 0 |
| Rest post | 123 ± 8 | 120 ± 0 |
| Task 1 | 240 ± 105 | 301 ± 148 |
| Task 2 | 250 ± 146 | 377 ± 245 |
| Task 3 | 132 ± 47 | 433 ± 187 |
| Task 4 | 484 ± 200 | 448 ± 328 |
| Instructions | 870 ± 238 | 403 ± 64 |
| Feedback | 554 ± 151 | 887 ± 160 |

### Preprocessing and analysis of ECG data

Raw ECG signal for each participant was extracted from the original data set as an EDF file using EDF Browser and imported into Kubios HRV Premium software (version 3.5.0, Biosignal Analysis and Medical Imaging Group, University of Eastern Finland, Kuopio, Finland). First, the ECG data was preprocessed using automatic beat correction (5% acceptance threshold). Thereafter the remaining artifacts, such as missing or incorrectly detected R peaks of the QRS complex, were visually inspected and manually corrected if needed. Moreover, noisy data segments were marked as noise and removed from further analyses.

Finally, both time-domain and frequency-domain measures of HRV were computed from the detrended (smoothing priors method, smoothing parameter 500, cutoff frequency 0.035 Hz) and interpolated (equidistant sampling, interpolation rate 4 Hz) RR interval series. Time-domain measures included mean interval between successive RR peaks (mean RR), standard deviation of all NN intervals (SDNN), the root-mean square of the differences of successive RR intervals (RMSSD), number of pairs of adjacent NN intervals differing by more than 50 ms (NN50), NN50 divided by the total number of all NN intervals (pNN50) and mean, minimum and maximum HR. Frequency-domain measures were extracted from spectra estimated using Fast Fourier Transform (FFT; Welch’s periodogram method, window width 300 s, window overlap 50%). Frequency-domain measures (i.e., peak frequency and power) were calculated separately for each frequency band: very low frequency (0-0.04 Hz), low frequency (0.4-0.15 Hz) and high frequency (0.15-0.4 Hz) (Acharya et al., 2006; Quigley et al., 2024; Tarvainen et al., 2014).

In the further analyses, we focused on well-established measures of cardiac activity, mean HR and RMSSD, which reflect the physiological state of the body (Berntson et al., 1997; TFESCNASP Electrophysiology, 1996; Quigley et al., 2024). While heart rate is generally considered an indicator of the combined influences of the sympathetic and parasympathetic branches of the autonomic nervous system (Acharya et al., 2006; Sacha, 2014), RMSSD is regarded as an indicator of parasympathetic activity (Acharya et al., 2006; Bertsch et al., 2012). Both measures are widely used in research and clinical practice, and are highly applicable in both laboratory settings and diverse everyday life contexts.

Although some time- and frequency-domain measures of HRV, such as RMSSD and HF-HRV, are often considered comparable, we focus on RMSSD because it can be reliably applied to short-term recordings and appears to be less affected by variations in respiratory parameters than its frequency-domain counterpart, HF-HRV (e.g., Quigley et al., 2024; Penttilä et al., 2001; but see Grossman & Kollai, 1993; Ritz, 2024). This consideration is particularly important in the present study, as the students and instructors interacted during the naturalistic learning situation, where respiratory parameters likely varied due to speaking.

### Preprocessing and analysis of EEG data

Raw EEG signal for each participant was extracted from the original data set as an EDF file that was then converted into a FIF file. Bad channels were detected and interpolated using PyPREP library (Bigdley-Shamlo et al., 2015). Any remaining bad channels were excluded in the forthcoming analysis steps. MNE-Python library version 0.24.0 (Gramfort et al., 2013) was used to band-pass filter the EEG signals at 1-40 Hz using a zero phase 10 second Hamming window with a Fast Fourier Transform (FFT) method and a Finite Impulse Response (FIR) filter design. Lower and upper transition bandwidth was set to 0.5 Hz.

Independent component analysis (ICA) was used to remove artifacts of cardiac and ocular origin from the raw data. ICA was conducted using fastica algorithm (Hyvärinen, 1999; Hyvärinen & Oja, 2000) with 0.98 explained variance and maximum 2000 iterations. Electro-oculogram (EOG) channels were constructed from EEG channels (AF3 and AF4 for forestry; virtual bipolar reference channels FP1-AF3, FP2-AF4, F7-T7, F8-T8 for aviation) and bandpass filtered at 1-10 Hz. Channels included in virtual channels were only used as EOG channels and were thus removed from the further EEG analyses. Together with visual inspection, components that had a correlation above 0.4 with EOG channels were removed. To remove cardiac artifacts, ICA was fitted again on the raw data with ocular artifacts removed using the same parameters as in the previous step. The ECG channel was bandpass filtered at 8-16 Hz. ICA components related to cardiac activity were detected using cross-trial phase statistics and the z-score method with the default MNE-Python ICA objects’ find_bads_ecg method parameters.

After preprocessing, power spectral density of the EEG signals at 1-40 Hz was calculated using Welch’s method (length of the window 2048 samples, 50 % overlap, Hann window) resulting in a frequency resolution of 0.24 Hz. Fitting oscillations and one over f algorithm (FOOOF; Donoghue et al., 2020) was used to estimate the periodic (i.e., power and peak frequency of alpha-band oscillatory activity) and aperiodic components (i.e., exponent) of the oscillatory brain activity. Fitting was conducted on the channels F3, Fz, F4, C3, C1, Cz, C2, C4, P3, Pz, and P4 for the 64-channel montage and on channels F3, F4, C3, C4, P3, and P4 for the 13-channel montage. The frequency-band of interest was set to 7-14 Hz, maximum number of peaks to 8, peak width to 0.5-6 Hz, peak threshold to 2.0, minimum peak height to 0 and aperiodic mode to fixed. The mean power within alpha frequency band (7 – 14 Hz) was estimated from the flattened spectra that was calculated by subtracting the aperiodic activity from the original spectra.

Each FOOOF fit was visually inspected to verify the accuracy and correctness of the fit. FOOOF was fitted twice, 1) to detect the optimal fitting range at 1-40 Hz for every channel grouped by context, subject type, subject id, and segment label by inspecting the linearity of the power spectrum in log-log space and 2) to calculate the periodic and aperiodic components using the optimized fitting range. Log-log power spectrum linearity was defined by excluding lower 1 Hz to 3 Hz and upper 25 Hz to 40 Hz frequency ranges containing plateaus. If no lower and/or upper range plateaus were detected, the whole frequency range from 1 Hz to 40 Hz was used to fit FOOOF. Based on this inspection, we excluded the frequency ranges containing plateaus and used a fixed frequency range for all participants and segments during the calculation of periodic and aperiodic components.

Fitting range optimization was crucial to improve the detection of the peak center frequency in the alpha band. FOOOF failed to determine the alpha peak frequency when using the predetermined fitting range of 1–40 Hz in 26.8 % of cases within the aviation context and 27.3 % within the forestry context. In comparison, application of an optimized fitting range reduced the failure rates to 16.1 % and 20.8 % in the aviation and forestry contexts, respectively.

Although the use of an optimized frequency range in the FOOOF analysis improved alpha peak frequency detection, the algorithm still frequently failed to identify the peak accurately and reliably, particularly during task-related segments. Therefore, subsequent analyses focused on mean alpha power and exponent, which were more consistently and reliably detectable across segments.

### Statistical analysis

Statistical analyses were performed using IBM SPSS Statistics 30.0 (IBM Corp., Armonk, New York, USA). According to the Kolmogorov-Smirnov test not all measures of cardiac and brain activity were normally distributed. Thus, given the non-normality and the small sample size, a non-parametric Wilcoxon Signed-Rank test was performed to investigate the differences in cardiac and brain activity across the different phases of the naturalistic learning situation and across tasks varying in difficulty. To reduce the likelihood of false positives (i.e., type I error), a Bonferroni-adjusted significance threshold was applied, resulting in p < 0.007 when examining the differences in cardiac and brain activity across the phases of the naturalistic learning situation, and p < 0.008 when examining the differences in cardiac and brain activity between tasks of varying difficulty.

## RESULTS

Descriptive statistics of cardiac and brain activity are reported in Supplementary Information. Across all phases of the naturalistic learning situation, measures of cardiac activity fell within the normal reference range (Nunan et al., 2010; Shaffer & Ginsberg, 2017). Visual inspection of the cardiac activity (Figure 3) revealed that students’ heart rate was lowest during the rest before and after the simulation, increased during the instruction and feedback phases, and was highest during the task phases. As expected, the opposite pattern was observed for heart rate variability, reflecting the typical inverse relationship between mean HR and RMSSD. In instructors, heart rate and heart rate variability followed a similar trend to that of students, but heart rate variability peaked during the instruction and feedback phases.

**Figure 3.**
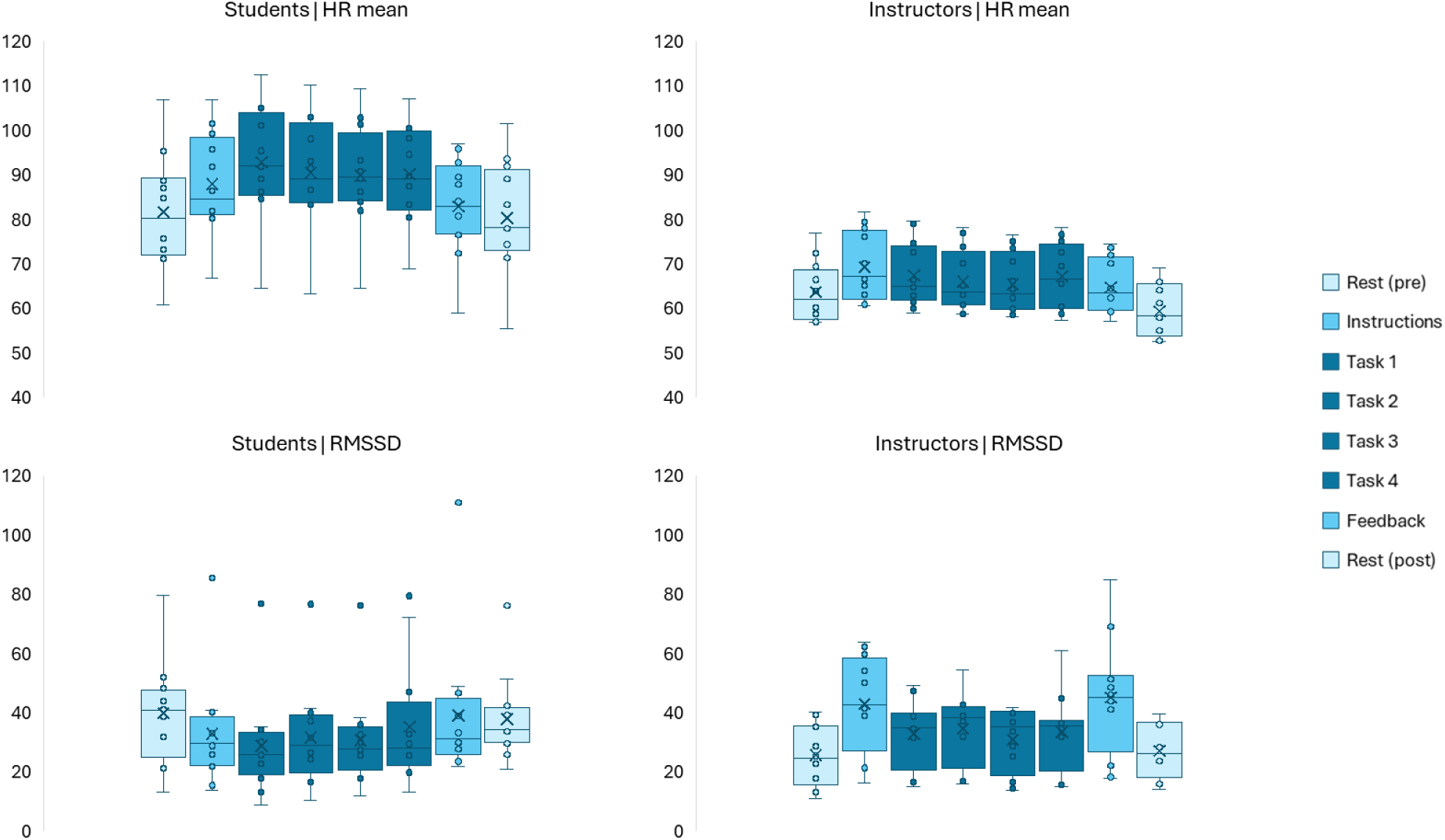
Cardiac activity across the phases of the naturalistic learning situation. In the box plots the upper and lower box boundaries represent the 75th and 25th percentiles, respectively, whiskers denote minimum and maximum values, cross the mean value, line inside the box the median value, and circles outside the whiskers are considered outliers.

Visual inspection of the brain activity indicated that alpha power was highest during the rest before and after the simulation, decreased during the instruction and feedback phases, and was lowest during the tasks in both students and instructors (Figure 4). No clear pattern was observed for the exponent (Figure 5).

**Figure 4.**
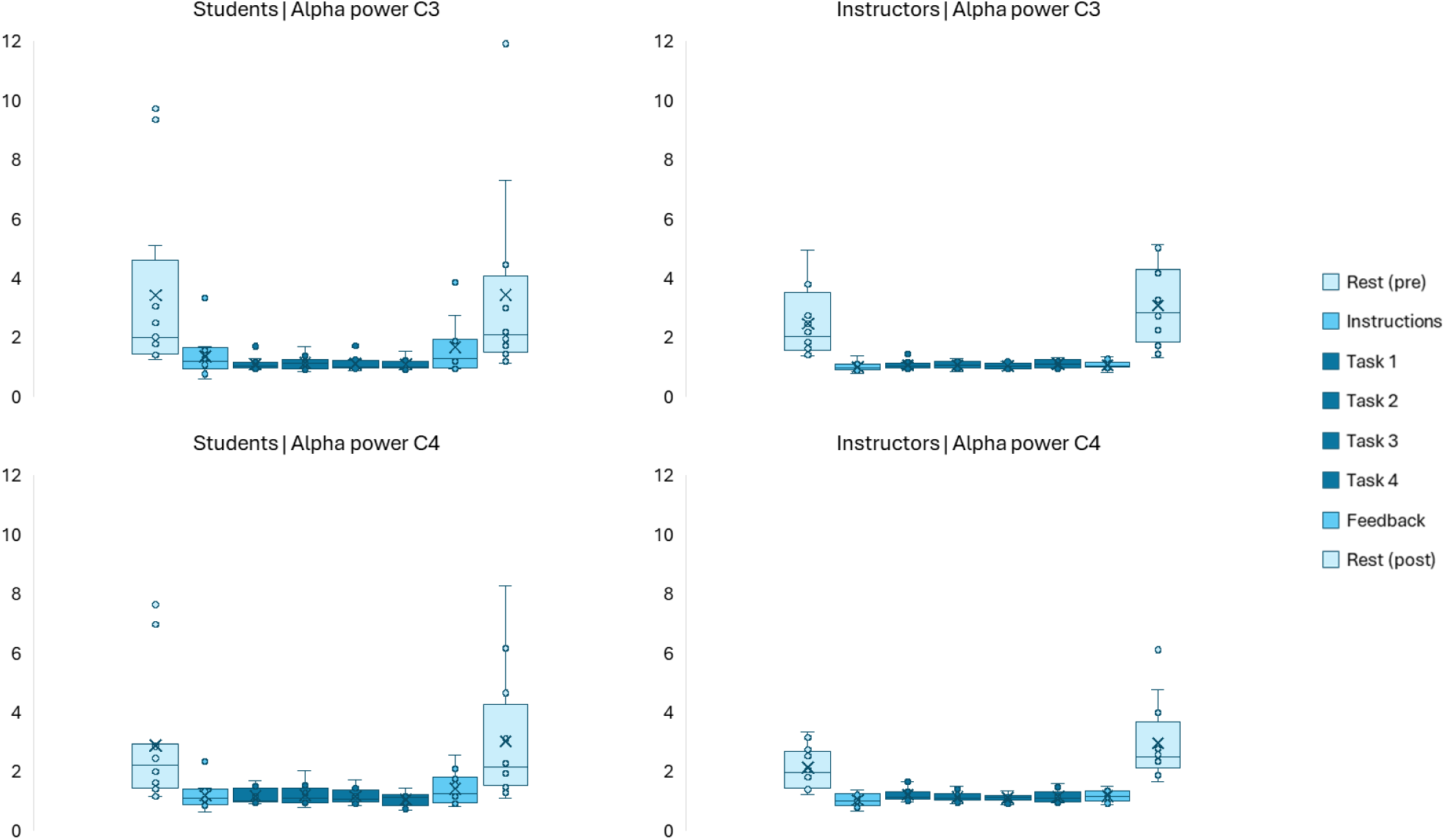
Alpha power across the phases of the naturalistic learning situation. In the box plots the upper and lower box boundaries represent the 75th and 25th percentiles, respectively, whiskers denote minimum and maximum values, cross the mean value, line inside the box the median value, and circles outside the whiskers are considered outliers.

**Figure 5.**
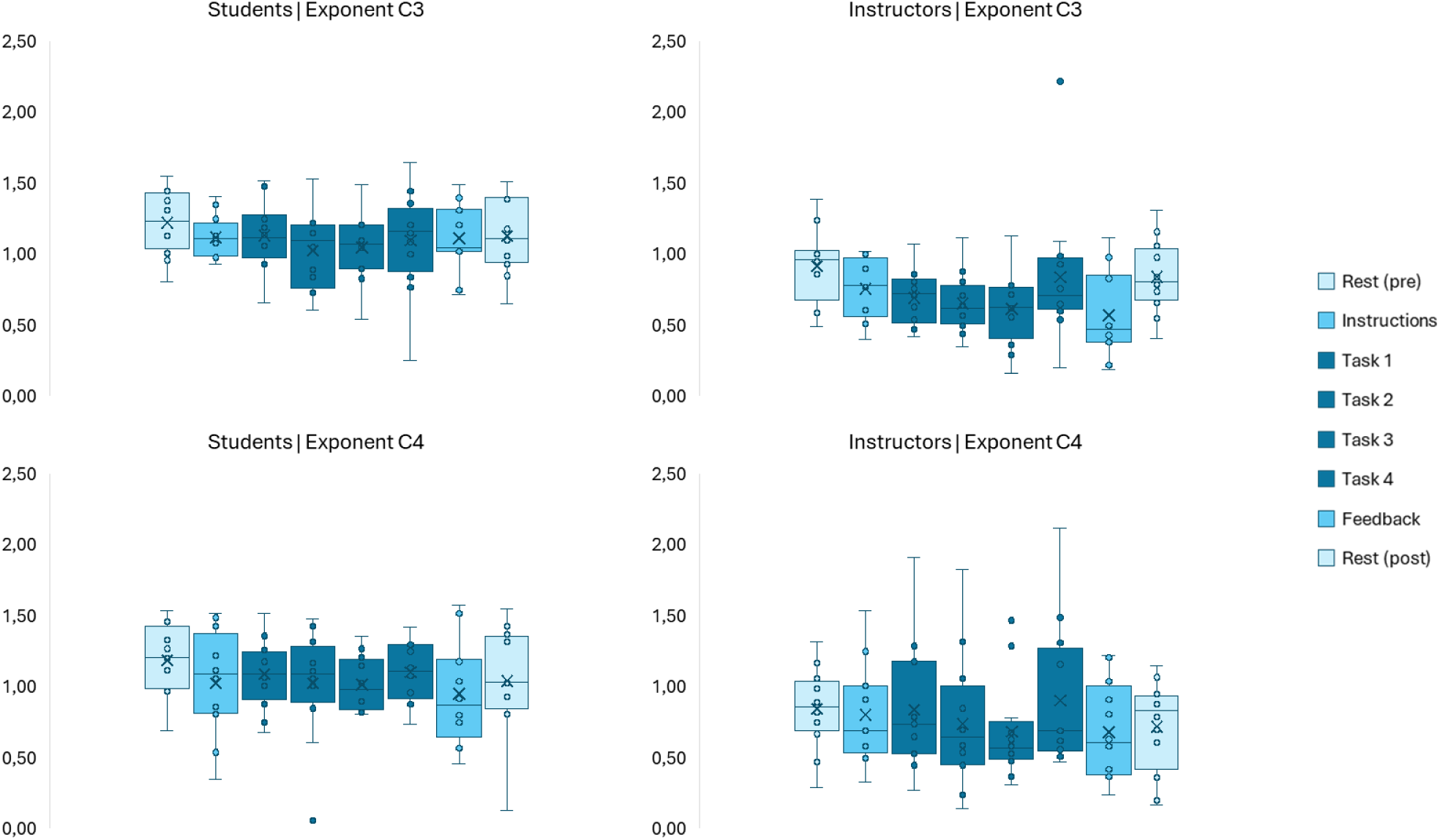
Exponent across the phases of the naturalistic learning situation. In the box plots the upper and lower box boundaries represent the 75th and 25th percentiles, respectively, whiskers denote minimum and maximum values, cross the mean value, line inside the box the median value, and circles outside the whiskers are considered outliers.

### Cardiac activity

#### Students

The Wilcoxon Signed-Rank test results indicated that students’ mean HR was lower during rest pre compared to task 1 (Z = −2.981, p = 0.003), task 2 (Z = −2.746, p = 0.006), and instructions (Z = −3.059, p = 0.002) phases (Table 2). No statistically significant differences in mean HR between tasks varying in difficulty were observed.

**Table 2.** Differences in students’ cardiac and brain activity across different phases of the naturalistic learning situation (rest pre vs. other phases) (Bonferroni-adjusted p < 0.007).

|  | Task 1 | Task 2 | Task 3 | Task 4 | Instructions | Feedback | Rest post |
| --- | --- | --- | --- | --- | --- | --- | --- |
| HR mean | 0.003** | 0.006** | 0.012 | 0.019 | 0.002** | 0.480 | 0.530 |
| RMSSD | 0.015 | 0.041 | 0.028 | 0.099 | 0.041 | 0.638 | 0.308 |
| Alpha power C3 | 0.002** | 0.002** | 0.002** | 0.003** | 0.002** | 0.004** | 1.000 |
| Alpha power C4 | 0.002** | 0.003** | 0.002** | 0.003** | 0.003** | 0.005** | 0.695 |
| Exponent C3 | 0.059 | 0.011 | 0.011 | 0.170 | 0.034 | 0.041 | 0.106 |
| Exponent C4 | 0.307 | 0.224 | 0.065 | 0.195 | 0.117 | 0.028 | 0.117 |

No statistically significant differences in RMSSD across the different phases of the naturalistic learning situation were observed. However, students’ RMSSD was statistically significantly lower during task 1 in comparison with task 2 (Z = −2.903, p = 0.004).

#### Instructors

Instructors’ mean HR was lower during rest pre in comparison with task 1 (Z = −3.059, p = 0.002) and instructions phases (Z = −3.059, p = 0.002), whereas mean HR during rest pre was higher than during rest post (Z = −2.903, p = 0.004) phase (Table 3). Moreover, mean HR was statistically significantly higher during task 1 compared to task 2 (Z = −2.981, p = 0.003).

**Table 3.** Differences in instructors’ cardiac and brain activity across different phases of the naturalistic learning situation (rest pre vs. other phases) (Bonferroni-adjusted p < 0.007).

|  | Task 1 | Task 2 | Task 3 | Task 4 | Instructions | Feedback | Rest post |
| --- | --- | --- | --- | --- | --- | --- | --- |
| HR mean | 0.002** | 0.012 | 0.209 | 0.041 | 0.002** | 0.209 | 0.004** |
| RMSSD | 0.015 | 0.015 | 0.117 | 0.008 | 0.002** | 0.002** | 0.308 |
| Alpha power C3 | 0.002** | 0.002** | 0.002** | 0.002** | 0.002** | 0.002** | 0.092 |
| Alpha power C4 | 0.006** | 0.002** | 0.002** | 0.002** | 0.002** | 0.003** | 0.071 |
| Exponent C3 | 0.091 | 0.041 | 0.041 | 0.480 | 0.077 | 0.031 | 0.326 |
| Exponent C4 | 0.859 | 0.456 | 0.158 | 0.814 | 0.583 | 0.182 | 0.272 |

Instructors’ RMSSD was statistically significantly lower during rest pre compared to instructions (Z = −-3.059, p = 0.002) and feedback phases (Z = −3.059, p = 0.002). No statistically significant differences in RMSSD between tasks were observed.

### Brain activity

#### Students

Statistically significant differences were observed in students’ alpha power between rest pre and all the other phases of the naturalistic learning situation except for rest post phase (Table 2). Specifically, students’ alpha power in sensor C3 was higher during rest pre in comparison to task 1 (Z = −3.059, p = 0.002), task 2 (Z = −3.059, p = 0.002), task 3 (Z = −3.059, p = 0.002), task 4 (Z = −2.981, p = 0.003), instructions (Z = −3.061, p = 0.002), and feedback phases (Z = −2.903, p = 0.004). Similarly, students’ alpha power in sensor C4 was higher during rest pre in comparison to task 1 (Z = −3.059, p = 0.002), task 2 (Z = −2.981, p = 0.003), task 3 (Z = −3.061, p = 0.002), task 4 (Z = −2.981, p = 0.003), instructions (Z = −2.943, p = 0.003), and feedback phases (Z = −2.803, p = 0.005). No statistically significant differences in alpha power between tasks were observed.

No statistically significant differences in exponent across the different phases of the naturalistic learning situation or between tasks were observed.

#### Instructors

Statistically significant differences were observed in instructors’ alpha power between rest pre and all the other phases of the naturalistic learning situation except for rest post phase (Table 3). Instructors’ alpha power in sensor C3 was higher during rest pre in comparison to task 1 (Z = −3.062, p = 0.002), task 2 (Z = −3.059, p = 0.002), task 3 (Z = −3.059, p = 0.002), task 4 (Z = −3.061, p = 0.002), instructions (Z = −3.059, p = 0.002), and feedback phases (Z = −3.061, p = 0.002). Similarly, instructors’ alpha power in sensor C4 was higher during rest pre in comparison to task 1 (Z = −2.756, p = 0.006), task 2 (Z = −3.059, p = 0.002), task 3 (Z = −3.059, p = 0.002), task 4 (Z = −3.061, p = 0.002), instructions (Z = −3.061, p = 0.002), and feedback phases (Z = −2.981, p = 0.003). No statistically significant differences in alpha power between tasks varying in difficulty were observed.

No statistically significant differences in exponent across the different phases of the naturalistic learning situation were observed. However, exponent in sensor C4 was lower during task 3 than during task 4 (Z = −2.677, p = 0.007).

### Qualitative examination of the reproducibility of cardiac and brain activity across repeated measurements

Since each instructor participated in the study three times with different students, this experimental design also allowed us to explore the reproducibility of cardiac and brain measures across the repetitions. Reproducibility of cardiac and brain activity across the instructors’ three repeated measurements with different students was assessed by visual inspection of the data (Figure 6).

**Figure 6.**
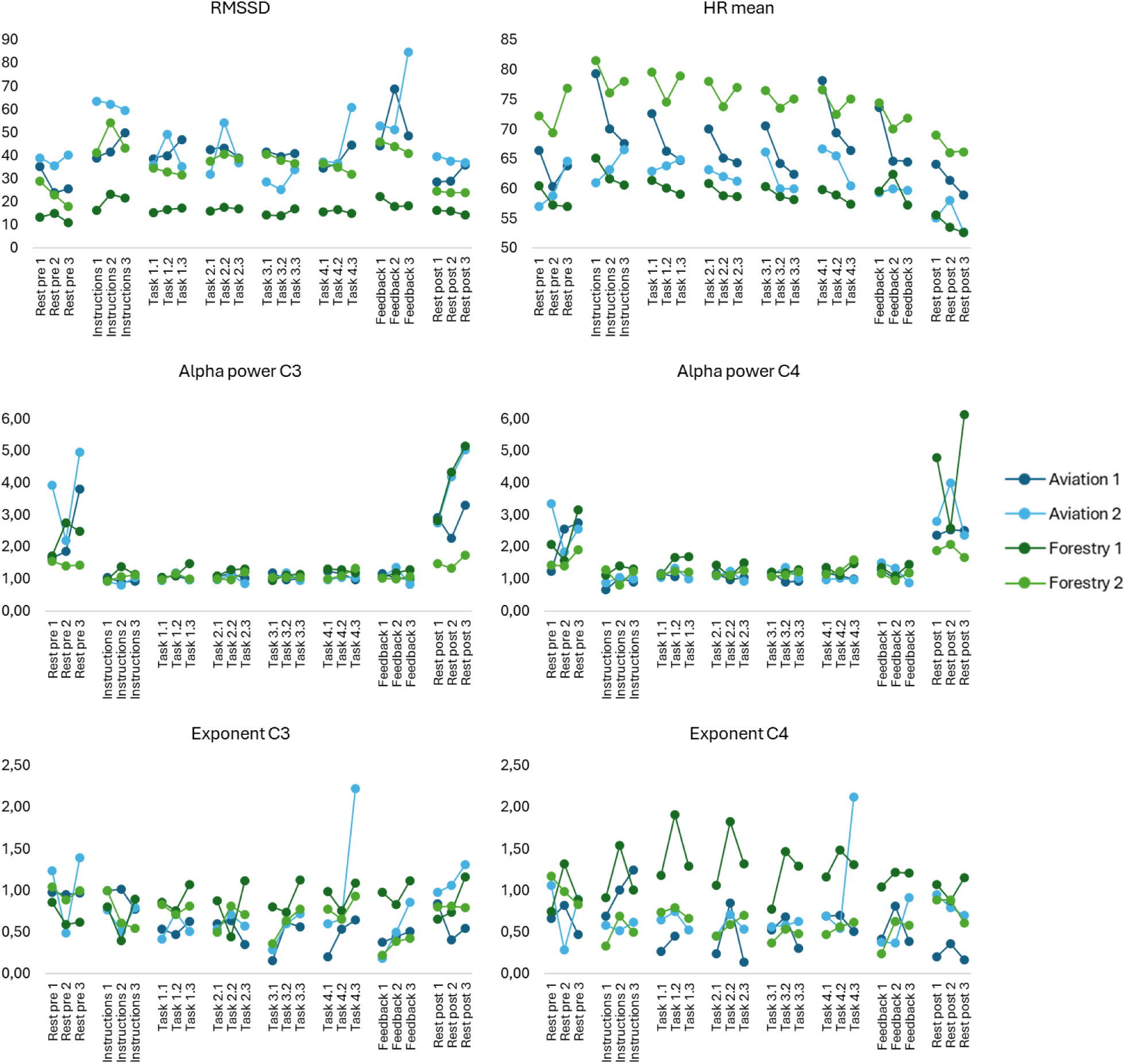
Reproducibility of instructors’ cardiac and brain activity across the different phases of the naturalistic learning situation. Each line represents one instructor (light and dark blue: aviation; light and dark green: forestry).

Visual inspection of the instructors’ cardiac activity indicated that, although baseline levels of heart rate and heart rate variability differed across instructors, these measures remained stable across repetitions. Visual inspection of alpha power showed that brain activity during rest varied across instructors and repetitions. However, during task-related phases of the naturalistic learning situation all instructors showed a comparable brain activity pattern both across instructors and repetitions. The exponent followed a similar pattern to the cardiac activity, indicating differences in the baseline levels across instructors yet demonstrating inconsistent responses across repetitions. Overall, measures of cardiac activity appeared stable across repetitions, whereas measures of brain activity were somewhat less consistent.

## DISCUSSION

This study investigated students’ and instructors’ cardiac and brain activity across the progression of the naturalistic learning situations in aviation and forestry contexts. First aim of this study was to capture robust and reliable features of cardiac and brain activity that reflect learning experiences during naturalistic learning situations. Yet, conducting neurophysiological recordings in naturalistic contexts with a level of quality sufficient to compute validated markers of known brain states, such as rhythmic brain activity across different frequency bands, is not trivial. Despite this challenge, we were able to successfully measure and extract features from both cardiac and brain activity that were associated with behaviourally and pedagogically meaningful phases during the naturalistic learning situation. However, this required a carefully designed experimental setup, ensuring data quality during data acquisition, as well as comprehensive preprocessing, artifact correction, and analysis of the data using standardized procedures. The most important step from the perspective of analysis, was structuring the naturalistic situation (i.e., the timeline of events) to enable the examination of the cardiac and brain measures with the required accuracy. This improved the signal-to-noise ratio to the extent that allowed accurate and reliable estimation of cardiac and brain activity across the phases of the naturalistic learning situation.

Importantly, the students’ and instructors’ cardiac and brain activity varied in a systematic manner along the phases of the naturalistic learning situation and across tasks with different levels of difficulty. In accordance with previous literature (e.g., Fairclough & Venables, 2006; Hughes et al., 2019), students’ heart rate was higher during instructions and first two tasks in comparison with the rest before the naturalistic learning situation. This observation of higher heart rate during task-related activities likely reflects task engagement and facilitation of optimal vigilance state for learning (Pendleton et al., 2016; Siennicka et al., 2019). Although previous studies have observed that heart rate responds to increasing task demands and cognitive load (Kahneman et al., 1969; Koskelo et al., 2024; Luque-Casado et al., 2016; Mansikka et al., 2016), after the first two tasks heart rate seemed to decrease closer to rest levels as there were no differences between rest and the last two tasks or between tasks varying in difficulty. This may be explained by the fact that these students are still novice and thus may react more strongly to the start of the learning situation rather than showing variability in their physiological responses across the varying task demands and difficulty levels.

We observed no differences in students’ heart rate variability across the different phases of the naturalistic learning situation. However, lower heart rate variability was observed during the first task in comparison with the second task suggesting that parasympathetic activity is reduced in the beginning of the active phases of the naturalistic learning situation. Together these findings suggest that cardiac activity varies across the phases of the naturalistic learning situation and to some extent also show sensitivity to task-related demands.

Examination of the students’ brain activity showed that alpha power was higher during rest before the naturalistic learning situation in comparison with all the other phases except for the rest after the naturalistic learning situation. This finding shows a pattern of higher alpha power during wakeful rest and lower alpha power during task-related phases, that is, states of active information processing, replicating the widely reported alpha power modulations in the literature (Foxe & Snyder, 2011; Jensen & Mazaheri, 2010; Klimesch, 2012). However, although previous studies have shown that the strength of alpha-band activity dynamically responds to the changes in cognitive load and task difficulty (Chikhi et al., 2022; Haegens et al., 2014), we did not observe differences in alpha power between tasks with varying difficulty. The lack of dynamic changes within alpha power across the tasks may be attributed to the use of highly immersive simulation environments, where visually engaging stimuli likely induced strong modulations in alpha power related to visual processing, thus reducing the task specific differences reflecting variations in task difficulty. On the other hand, for novice students all the tasks may remain challenging, requiring greater effort than for experts and more experienced learners.

Similar to students, also instructors showed higher heart rate during instructions and the first task in comparison with the rest before the naturalistic learning situation. Moreover, when the rest phases before and after the naturalistic learning situation were compared, lower heart rate was observed afterwards indicating relaxation and recovery after completing the naturalistic learning situation. Instructors’ heart rate was also higher during the first task compared to the second task indicating increased vigilance in the beginning of the active phases of the naturalistic learning situation. Against our expectations, lower heart rate variability was observed during rest before the naturalistic learning situation in comparison with the instructions and feedback. However, this finding needs to be interpreted with caution, as respiratory parameters have been shown to influence measures of heart rate variability (Grossman & Kollai, 1993; Quigley et al., 2024; Ritz, 2024). Since the instructors were actively speaking during the instructions and feedback, the observed differences may primarily reflect differences in respiratory activity rather than changes in the functional state of the autonomic nervous system. To address this in the future studies, including measures of respiratory changes would increase the interpretability of the results.

The comparable cardiac and activity patterns observed in instructors and students suggest that the autonomic and central nervous system resources are similarly utilized by both the instructor and the student, at least in interactive technological learning situations. In comparison to rest before and after the learning situation, alpha power was lower during the active task phases. Although the exponent of the aperiodic activity did not differ across the phases of the naturalistic learning situation, it was sensitive to task difficulty, being lower during the third task compared to the fourth task. Previous simulation and experimental studies utilizing measures of aperiodic activity, specifically the steepness of the aperiodic slope, have convincingly suggested that it reflects the level of inhibitory vs. excitatory neurotransmission at the system level (Donoghue et al., 2020; He, 2014). Interpreted in this context, the steeper aperiodic slope observed during the fourth task would indicate stronger inhibitory activity.

When considering individual variability, we observed that, despite inter-individual differences in baseline levels of cardiac and brain activity, all students and instructors showed similar variation of cardiac and brain activity across the different phases of the naturalistic learning situation. This suggests that, in addition to trait-like individual differences in cardiac and brain activity, these measures also fluctuate in a state-like manner across pedagogically and behaviourally meaningful phases of the learning process.

On the same note, based on visual inspection, cardiac measures appeared more consistent across repetitions, whereas the level of alpha activity was less reproducible. Thus, this finding suggests that some indicators of the functional state of the body and the brain are more stable across repeated measurements possibly reflecting trait variation, whereas others may be more sensitive to state variation showing fluctuations across repeated measurements. Although further investigation with larger samples is warranted, our findings provide support to the idea that measures of cardiac and brain activity could be used as – perhaps complementary – indicators of meaningful state and trait variation within and across individuals when investigating individual variability in learning and teaching.

Altogether, our findings concerning the students’ and instructors’ cardiac and brain activity during the naturalistic learning situation emphasize the embodied nature of learning. Specifically, the observation that students’ and instructors’ cardiac and brain activity show similar patterns of reactivity across the phases of the naturalistic learning situation suggests that both students and instructors are engaged in the naturalistic learning situations not only at the behavioural level but also at the physiological and neural levels. This finding highlights the importance of advancing our understanding of the embodied aspects of both learning and teaching.

It is important to note that we cannot directly attribute the changes in bodily physiology or brain activity to specific states or processes, such as cognitive load, task engagement, attention, or emotions because from the perspective of neurophysiology, these different psychological and cognitive domains largely overlap. In other words, increased cognitive load, emotional engagement, and higher attentional demands may all increase heart rate, decrease heart rate variability, and decrease alpha power, albeit with potential differences in more fine-grained characteristics. Thus, to improve the interpretability of the results, self-reported experiences related to the learning process should be included (Silvennoinen et al., 2022). On the same note, it is not possible to disentangle whether the changes in cardiac and brain activity across the phases of the naturalistic learning situation reflect responses to task demands or the extent to which they are driven by self-regulation or co-regulation between instructors and students. Future studies should investigate these aspects to better understand how individual and dyadic- or group-level regulatory processes influence the functional state of the body and the brain during learning. However, since these processes are reciprocal, it is challenging to capture the causal influences in naturalistic contexts.

Since recent neuroscientific research has demonstrated that bodily functions are intertwined with rhythmic brain activity across different cortical areas (Candia-Rivera et al., 2024; Karjalainen et al., 2025; Kluger & Gross, 2021; Tsakiris & Critchley, 2016), it is reasonable to assume that individual differences observed in oscillatory brain activity may at least partially reflect differences in the body-brain interaction (Azzalini et al., 2019). Moreover, recent studies have indicated that interactions between the physiological state of the body and brain activity are also linked to cognitive and affective processes as well as behavior (Criscuolo et al., 2022; Forte & Casagrande, 2025; Parviainen et al., 2022; Varga & Heck, 2017; Waselius et al., 2022). Therefore, future studies conducted in naturalistic settings should also examine the variation in physiological and brain signalling to improve our understanding of whether and how the body and the brain are intertwined during learning. This would also allow the examination of the directionality of the interactions, namely to what extent the interactions reflect top-down (i.e., brain-body) and bottom-up processes (i.e., body-brain).

Not only the interaction between body and brain signalling, that is, intra-individual body–brain coupling, is informative for understanding the learning experience, but also the inter-individual coupling, that is, the interaction between the student and the instructor. Specifically, investigating synchrony in cardiac or neural activity within a dyad (i.e., student and instructor) could provide novel insights into the physiological and neural mechanisms underlying interactive learning and collaboration. Previous studies have demonstrated that teacher-student brain-to-brain synchrony reflects the level of student engagement (Bevilacqua et al., 2019), and that brain-to-brain synchrony between students predicts social dynamics and engagement in classroom settings (Dikker et al., 2017). Thus, integrating measures of body-brain interaction within individuals with inter-individual synchrony would enable a multi-level investigation of physiological and neural mechanisms supporting learning-related processes.

Neuroimaging studies are typically low in ecological validity making it difficult to translate laboratory findings on learning to real-life contexts (van Atteveldt et al., 2018). At the same time, studies using neuroimaging methods in naturalistic learning situations are still rare. To support pedagogical insight, however, it is not enough to ensure high quality recordings in naturalistic contexts, but the recordings also need to be done in a way that they provide meaningful information from the pedagogical point of view. Thus, a key advantage of our study was capturing neurophysiological aspects of authentic learning situations that are integrated into the regular curriculum and teaching practices of the collaborating institutions. The strength of our study also lies in the inclusion of representatives of education sciences, education practitioners, and neuroscientists at all stages of the research project. Specifically, instructors contributed to the planning of the experimental design to ensure realistic and ecologically valid setup, as well as to the selection of the tasks that accurately reflected real-life demands and circumstances.

To systematically examine the physiological and neural states that optimally facilitate positive and supportive learning experience in naturalistic learning situations, it is necessary to ensure the high quality and unbiased interpretation of the data. This calls for standardized protocols to avoid misleading, biased, or overstated conclusion derived from limited evidence. In Table 4, we highlight key aspects that could help harmonize experimental settings for investigating intra- and inter-individual differences in the body and the brain in naturalistic learning situations.

**Table 4.** Key aspects for harmonizing experimental settings to investigate the body and the brain in naturalistic contexts.

| Experimental design and data acquisition | Data preprocessing and analysis | Interpretability of the findings |
| --- | --- | --- |
| • To increase cross-study interpretation of studies conducted in naturalistic contexts, efforts should be made, within practical limits and without compromising the authenticity, to control the experimental settings or alternatively a detailed description of the setup is recommended.<br>• Detailed reporting of the selection process with justification and the recording of biosignals is required to improve the replicability of the findings.<br>• When combining various biosignals, it is crucial to acknowledge data modality specific restrictions, such as optimal sampling rate, artifacts, and varying temporal scales of physiological and neural activity.<br>• To facilitate the examination of the interplay across different data modalities, temporal alignment of each modality already during data acquisition is crucial for subsequent data integration.<br>• Since data quality is typically poorer when recorded in naturalistic contexts (e.g., absence of electromagnetic shielding, natural movements, many sources of noise and artifacts) and the quality of the raw data affects the accuracy of extracting robust and reliable features from the recorded biosignals, ensuring data quality during data acquisition should be prioritized. Moreover, detailed documentation of the events during the investigation is necessary for the exclusion of intervening factors (e.g., artifacts related to speech and movement) and accurate interpretation of the biosignals. | • To enable quality control and replicability of studies, the use of existing guidelines for conducting and reporting physiological and neuroimaging research (e.g., Boucsein et al., 2012; Quigley et al., 2024; Urigüen & Garcia-Zapirain, 2015; Yücel et al., 2021) is recommended.<br>• State-of-the-art analysis methods should be applied to minimize sources of noise not only during data acquisition but also during data preprocessing.<br>• To enable reliable feature extraction from biosignals, analyses focusing on comparisons or contrasts between baseline or control conditions and experimental conditions can further reduce the sources of noise, confounding factors, and inter-individual variability. Especially in naturalistic contexts, inclusion of a control condition (e.g., resting state) would be beneficial for the analysis of the data. | • In general, conducting experiments first in laboratory settings, then in laboratories using naturalistic tasks and stimuli, and finally in fully naturalistic contexts is crucial for assessing the replicability of findings across contexts with varying levels of ecological validity, thereby improving the generalization and interpretability of the results (Cisek & Green, 2024; Hari et al., 2015; Janssen et al., 2021).<br>• Alternatively, it is recommended to validate the findings of previous laboratory studies in naturalistic contexts.<br>• Because interpreting results from studies conducted in naturalistic, less controlled settings is more challenging than in laboratory settings, it is recommended to focus on previously established features or measures and assess their replicability in naturalistic contexts. |

This study demonstrated that it is feasible to capture robust and reliable indicators of processes reflecting learning experience from cardiac and brain activity during naturalistic learning situations. Furthermore, the findings indicate that both students’ and instructors’ heart rate, heart rate variability, and characteristics of oscillatory brain activity vary systematically across behaviourally and pedagogically meaningful phases of the naturalistic learning situation. The findings of this study enhance our understanding of how body and brain activity vary during naturalistic learning situations, both within and across individuals, thereby shedding light on the nervous system determinants underlying individual differences in learning experience. Potentially, in the future, this information could also be applied at individual level to help individuals recognize how the functional state of their body and brain influences the learning process and develop self-regulation strategies to cope with learning-related emotions, such as stress, anxiety, or excitement. Indeed, it has been shown that learning interventions involving making brain activity visible and approachable to the students, enhances the learners’ agency and self-efficacy (Harris et al., 2021; Janssen & van Atteveldt, 2022). Neurophysiological measurements could, thus, be applied as tools to support self-regulation, assessment, and feedback in learning contexts. Studying naturalistic learning situations also allows examination of the body and the brain from the perspective of learning interactions, particularly how the functional state of the body and brain is linked to the dynamics of interaction between student and instructor. In the long term, neurophysiological measures could be used to develop tools that help students and instructors identify optimal learning states and to provide timely support. In this way, the study lays the groundwork for developing individually tailored learning paths that take into account individual nervous system level differences in learning as well as learners’ specific needs.

## ACKNOWLEDGEMENTS

This work was supported by European Regional Development Fund (React-EU, grant number A75070), Regional Council of Central Finland (grant number KSL/201/04.03.02.00/2018), and Research Council of Finland (grant number 365343). The funders had no role in study design, data collection and analysis, decision to publish, or preparation of the manuscript. The authors wish to express their thanks to the members of “Development of educational service ecosystem using physiological data and intelligent systems” project team as well as the students and instructors of the Finnish Air Force Academy and Poke Vocational College for collaboration.

## DATA AVAILABILITY

The data cannot be made openly available according to the ethical permission and national privacy regulations at the time of the study.

## CREDIT AUTHORSHIP CONTRIBUTION STATEMENT

Suvi Karjalainen: Writing – review & editing, Writing – original draft, Data curation, Methodology, Investigation, Formal analysis, Conceptualization

Juha Leukkunen: Writing – review & editing, Writing – original draft, Software, Data curation, Methodology, Formal analysis

Tiina Kullberg: Writing – review & editing, Data curation, Methodology, Investigation

Tiina Parviainen: Writing – review & editing, Supervision, Methodology, Funding acquisition, Conceptualization

