## Supplementary Information for "Body and brain signalling during naturalistic learning situations"

**SI1.** Descriptive statistics of cardiac and brain activity for students and instructors in both aviation and forestry simulation-based learning situations. Please note that for exponent at sensor C3 during task 3, n = 11 for all students and instructors, and n = 5 for aviation students and instructors.

| | | Students \| All | Students \| Aviation | Students \| Forestry | Instructors \| All | Instructors \| Aviation | Instructors \| Forestry |
|---|---|---|---|---|---|---|---|
| HR (beats / min) | Rest pre | 81.78 ± 12.67 | 73.57 ± 9.06 | 89.98 ± 10.46 | 63.76 ± 6.56 | 61.89 ± 3.76 | 65.63 ± 8.50 |
| | Rest post | 80.48 ± 12.36 | 74.27 ± 10.93 | 86.70 ± 11.15 | 59.47 ± 5.84 | 58.41 ± 4.13 | 60.54 ± 7.43 |
| | Task 1 | 92.96 ± 12.71 | 87.63 ± 15.41 | 98.29 ± 7.06 | 67.51 ± 7.21 | 65.97 ± 3.51 | 69.04 ± 9.82 |
| | Task 2 | 90.54 ± 12.43 | 86.99 ± 15.21 | 94.08 ± 8.86 | 66.17 ± 6.95 | 64.41 ± 3.18 | 67.92 ± 9.42 |
| | Task 3 | 89.99 ± 11.56 | 87.61 ± 14.39 | 92.36 ± 8.56 | 65.51 ± 6.80 | 63.94 ± 4.11 | 67.09 ± 8.89 |
| | Task 4 | 90.29 ± 10.81 | 88.46 ± 13.89 | 92.12 ± 7.47 | 67.36 ± 7.25 | 67.92 ± 5.86 | 66.80 ± 8.98 |
| | Instructions | 88.08 ± 11.33 | 81.86 ± 10.37 | 94.30 ± 9.07 | 69.34 ± 7.69 | 68.05 ± 6.51 | 70.63 ± 9.16 |
| | Feedback | 82.98 ± 10.85 | 79.26 ± 11.71 | 86.71 ± 9.42 | 64.86 ± 6.21 | 63.71 ± 5.51 | 66.01 ± 8.70 |
| RMSSD (ms) | Rest pre | 40.05 ± 17.23 | 47.46 ± 18.93 | 32.64 ± 12.78 | 25.85 ± 10.26 | 33.45 ± 6.97 | 18.25 ± 6.66 |
| | Rest post | 38.01 ± 14.35 | 43.81 ± 17.98 | 32.21 ± 7.00 | 27.35 ± 9.01 | 34.80 ± 4.78 | 19.90 ± 4.73 |
| | Task 1 | 29.04 ± 17.11 | 35.41 ± 22.54 | 22.68 ± 6.21 | 33.02 ± 11.32 | 41.25 ± 5.83 | 24.78 ± 9.23 |
| | Task 2 | 31.87 ± 17.03 | 38.30 ± 21.65 | 25.44 ± 8.37 | 34.83 ± 11.97 | 41.57 ± 7.59 | 28.10 ± 12.21 |
| | Task 3 | 30.90 ± 16.19 | 35.92 ± 21.72 | 25.88 ± 6.68 | 31.09 ± 10.88 | 35.22 ± 7.03 | 26.96 ± 13.05 |
| | Task 4 | 35.44 ± 20.65 | 45.35 ± 26.09 | 25.52 ± 4.65 | 33.62 ± 13.07 | 41.96 ± 10.02 | 25.28 ± 10.41 |
| | Instructions | 33.00 ± 18.67 | 40.58 ± 22.92 | 25.42 ± 10.17 | 43.21 ± 16.10 | 52.92 ± 10.77 | 33.50 ± 15.09 |
| | Feedback | 39.27 ± 24.16 | 49.36 ± 31.56 | 29.18 ± 6.59 | 45.25 ± 19.73 | 58.74 ± 15.41 | 31.76 ± 13.50 |
| Alpha power C3 | Rest pre | 3.44 ± 3.04 | 3.29 ± 3.04 | 3.60 ± 3.32 | 2.48 ± 1.16 | 3.07 ± 1.35 | 1.90 ± 0.58 |
| | Rest post | 3.46 ± 3.18 | 4.13 ± 3.97 | 2.78 ± 2.33 | 3.11 ± 1.33 | 3.41 ± 1.03 | 2.81 ± 1.61 |
| | Task 1 | 1.12 ± 0.71 | 1.22 ± 0.28 | 1.03 ± 0.05 | 1.08 ± 0.14 | 1.04 ± 0.09 | 1.13 ± 0.18 |
| | Task 2 | 1.17 ± 0.23 | 1.22 ± 0.31 | 1.12 ± 0.13 | 1.09 ± 0.13 | 1.04 ± 0.11 | 1.15 ± 0.14 |
| | Task 3 | 1.13 ± 0.22 | 1.22 ± 0.29 | 1.05 ± 0.06 | 1.06 ± 0.09 | 1.06 ± 0.11 | 1.06 ± 0.07 |
| | Task 4 | 1.11 ± 0.20 | 1.13 ± 0.24 | 1.09 ± 0.155 | 1.14 ± 0.14 | 1.07 ± 0.11 | 1.21 ± 0.14 |
| | Instructions | 1.38 ± 0.71 | 1.14 ± 0.44 | 1.62 ± 0.88 | 1.02 ± 0.15 | 0.94 ± 0.10 | 1.10 ± 0.17 |

| | | | | | | | |
|---|---|---|---|---|---|---|---|
| | Feedback | 1.69 ± 0.89 | 1.36 ± 0.48 | 2.01 ± 1.12 | 1.09 ± 0.14 | 1.07 ± 0.17 | 1.11 ± 0.11 |
| Alpha power C4 | Rest pre | 2.90 ± 2.17 | 2.98 ± 2.37 | 2.82 ± 2.18 | 2.16 ± 0.71 | 2.38 ± 0.74 | 1.93 ± 0.66 |
| | Rest post | 3.05 ± 2.22 | 3.55 ± 2.61 | 2.54 ± 1.84 | 2.98 ± 1.32 | 2.76 ± 0.63 | 3.19 ± 1.83 |
| | Task 1 | 1.20 ± 0.28 | 1.33 ± 0.34 | 1.07 ± 0.14 | 1.24 ± 0.23 | 1.12 ± 0.13 | 1.35 ± 0.26 |
| | Task 2 | 1.23 ± +0.35 | 1.32 ± 0.43 | 1.14 ± 0.26 | 1.17 ± 0.17 | 1.08 ± 0.11 | 1.26 ± 0.18 |
| | Task 3 | 1.18 ± 0.26 | 1.29 ± 0.33 | 1.08 ± 0.13 | 1.14 ± 0.13 | 1.10 ± 0.17 | 1.19 ± 0.07 |
| | Task 4 | 1.07 ± 0.22 | 1.07 ± 0.28 | 1.08 ± 0.18 | 1.18 ± 0.21 | 1.04 ± 0.09 | 1.32 ± 0.20 |
| | Instructions | 1.21 ± 0.44 | 1.05 ± 0.31 | 1.36 ± 0.52 | 1.06 ± 0.22 | 0.92 ± 0.14 | 1.20 ± 0.21 |
| | Feedback | 1.45 ± 0.54 | 1.23 ± 0.40 | 1.66 ± 0.60 | 1.20 ± 0.20 | 1.20 ± 0.23 | 1.20 ± 0.18 |
| Exponent C3 | Rest pre | 1.23 ± 0.23 | 1.14 ± 0.21 | 1.32 ± 0.22 | 0.92 ± 0.26 | 1.00 ± 0.31 | 0.83 ± 0.19 |
| | Rest post | 1.13 ± 0.26 | 1.16 ± 0.22 | 1.11 ± 0.32 | 0.84 ± 0.25 | 0.86 ± 0.33 | 0.83 ± 0.17 |
| | Task 1 | 1.14 ± 0.24 | 1.05 ± 0.22 | 1.22 ± 0.24 | 0.70 ± 0.19 | 0.55 ± 0.12 | 0.84 ± 0.12 |
| | Task 2 | 1.03 ± 0.27 | 1.02 ± 0.23 | 1.04 ± 0.33 | 0.66 ± 0.21 | 0.57 ± 0.12 | 0.74 ± 0.25 |
| | Task 3 | 1.05 ± 0.24 | 1.03 ± 0.28 | 1.07 ± 0.23 | 0.62 ± 0.26 | 0.49 ± 0.22 | 0.74 ± 0.25 |
| | Task 4 | 1.10 ± 0.36 | 1.03 ± 0.43 | 1.17 ± 0.30 | 0.84 ± 0.49 | 0.81 ± 0.71 | 0.87 ± 0.16 |
| | Instructions | 1.12 ± 0.15 | 1.07 ± 0.07 | 1.17 ± 0.19 | 0.76 ± 0.21 | 0.81 ± 0.19 | 0.71 ± 0.23 |
| | Feedback | 1.11 ± 0.24 | 1.07 ± 0.23 | 1.16 ± 0.27 | 0.57 ± 0.30 | 0.48 ± 0.22 | 0.66 ± 0.36 |
| Exponent C4 | Rest pre | 1.19 ± 0.25 | 1.16 ± 0.30 | 1.23 ± 0.21 | 0.85 ± 0.28 | 0.70 ± 0.28 | 0.99 ± 0.22 |
| | Rest post | 1.04 ± 0.37 | 1.08 ± 0.25 | 1,01 ± 0.49 | 0.72 ± 0.33 | 0.53 ± 0.33 | 0.92 ± 0.44 |
| | Task 1 | 1.09 ± 0.24 | 1.05 ± 0.22 | 1.13 ± 0.28 | 0.84 ± 0.46 | 0.53 ± 0.18 | 1.10 ± 0.47 |
| | Task 2 | 1.02 ± 0.39 | 1.04 ± 0.24 | 1.02 ± 0.52 | 0.74 ± 0.48 | 0.49 ± 0.27 | 0.99 ± 0.52 |
| | Task 3 | 1.02 ± 0.19 | 1.01 ± 0.18 | 1.03 ± 0.22 | 0.69 ± 0.35 | 0.55 ± 0.13 | 0.82 ± 0.46 |
| | Task 4 | 1.11 ± 0.22 | 1.14 ± 0.27 | 1.07 ± 0.16 | 0.91 ± 0.51 | 0.88 ± 0.61 | 0.94 ± 0.44 |
| | Instructions | 1.03 ± 0.37 | 1.06 ± 0.39 | 0.99 ± 038 | 0.80 ± 0.35 | 0.78 ± 0.29 | 0.83 ± 0.43 |
| | Feedback | 0.95 ± 0.36 | 0.96 ± 0.43 | 0.95 ± 0.31 | 0.68 ± 0.35 | 0.55 ± 0.25 | 0.82 ± 0.40 |
